# Analysis of the effectiveness of sub-sensory electrical noise stimulation during visuomotor adaptations in different visual feedback conditions

**DOI:** 10.1101/739722

**Authors:** Anna Margherita Castronovo, Ciara Giles Doran, Méabh Holden, Giacomo Severini

## Abstract

Sub-sensory electrical noise stimulation has been shown to improve motor performance in tasks that rely principally on proprioceptive feedback. During the generation of movements such as reaching, proprioceptive feedback combines dynamically with visual feedback. It is still unclear whether boosting proprioceptive information in tasks where proprioception mixes with vision can influence motor performance at all, either by improving or worsening it. To better understand this point, we tested the effect of electrical noise stimulation applied superficially to the muscle spindles during four different experiments consisting of isometric reaching tasks under different visual feedback conditions. The first experiment (n=40) consisted of a reach-and-hold task where subjects had to hold a cursor on a target for 30 seconds and had visual feedback removed 10 seconds into the task. Subjects performed 30 repetitions of this task with different stimulation levels, including no stimulation. We observed that trials in which the stimulation was present, displayed smaller movement variability. Moreover, we observed a positive correlation between the level of stimulation and task performance. The other experiments consisted of three versions of an isometric visuomotor adaptation task where subjects were asked to reach to random targets in less than 1.5 seconds (otherwise incurring in negative feedback) while overcoming a 45° clockwise rotation in the mapping between the force exerted and the movement of the cursor. The three experiments differed in the visual feedback presented to the subjects, with one group (n=20) performing the experiment with full visual feedback, one (n=10) with visual feedback restricted only to the beginning of the trajectory and one (n=10) without visual feedback of the trajectory. All subjects performed their experiment twice, with and without stimulation. We did not observe substantial effects of the stimulation when visual feedback was present (either completely or partially). We observed a limited effect of the stimulation in the absence of visual feedback consisting in a significant smaller number of negative-feedback trials in the first block of the adaptation phase. Our results suggest that sub-sensory stimulation can be beneficial when proprioception is the main feedback modality but mostly ineffective in tasks where visual feedback is actively employed.

## 1. Introduction

Mechanical and electrical noise stimulation targeting joints and muscles can alter the kinesthetic sense and lead to improved motor performances (Cordo et al., 1996; Gravelle et al., 2002; Priplata et al., 2002; Collins et al., 2003; Priplata et al., 2006; Ross and Guskiewicz, 2006; Mendez-Balbuena et al., 2012; Collins et al., 2014; Iliopoulos et al., 2014; Miranda et al., 2016; Severini and Delahunt, 2018). Mechanical noise stimulation directly modifies the response of sensory receptors, while electrical noise stimulation alters the baseline transmembrane potential of the stimulated afferents making them more likely to fire in response to a weak stimulus (Gravelle et al., 2002; Miranda et al., 2016). Both effects are supposedly related to stochastic resonance (SR), a phenomenon for which noise can improve the reception of weak signals in threshold-based systems (Gammaitoni, 1995). By the SR phenomenon, noise added to the input of a threshold-based receiving system can improve the detection of a weak input signal by spuriously amplifying it. Values of noise that are too low may not bring the weak signal above the receiving threshold, while values of noise that are too high risk to mask the characteristics of the input signal and thus lead to erroneous detections. Therefore, the SR phenomenon predicts the presence of an optimal level of stimulation that maximizes the performance of the receiving system.

The SR phenomenon has been observed to occur in response to noise stimulation in biological systems in general (Collins et al., 1995), and in human sensory receptors in particular (Cordo et al., 1996; Mendez-Balbuena et al., 2012; Iliopoulos et al., 2014; Mendez-Balbuena et al., 2015). Proprioception plays a crucial role during the execution and learning of voluntary movements (Fleishman and Rich, 1963; Sober and Sabes, 2003) and sensory deficits have been shown to affect motor re-learning after a neurological injury (Vidoni and Boyd, 2009). Several studies have shown that superficial electrical noise stimulation targeting sensory receptors at sub-sensorial current levels (intended as current levels that do not elicit conscious perception) can improve performance during different motor tasks in healthy subjects (Magalhaes and Kohn, 2012; Iliopoulos et al., 2014; Magalhaes and Kohn, 2014), elderlies (Gravelle et al., 2002) and individuals suffering from sensory loss (Collins et al., 2014). In all these experiments, the motor tasks selected (i.e. single leg stance) relied heavily on proprioception as sensory feedback modality. Recently, we were also able to show that, in opposition to the results obtained using sub-sensorial stimulation, supra-sensorial currents lead to a decrease in motor performance during mildly challenging balance tasks (Severini and Delahunt, 2018), although it is not clear whether this effect is caused by the conscious sensation or by the degradation in performance expected by the SR model for levels of noise above the optimal one. It has been proposed that sub-sensory noise stimulation could be used to improve the quality and quantity of available proprioceptive information during rehabilitation of patients affected by proprioceptive deficits (Collins et al., 2003). In this scenario, since motor learning in rehabilitation is often associated with complex tasks (e.g. walking, reaching…) where several sensory feedback modalities are integrated and employed at the same time, it is paramount to understand how boosting proprioception can affect the overall feedback information. This latter point is still unexplored in literature. In fact, while most studies employing sub-sensory stimulation have shown its benefits in tasks where proprioception is the main feedback modality, it is not clear what its effect would be in tasks where proprioception integrates (or competes) with other sensory modalities, such as visual. As a case in point, during reaching movements proprioceptive and visual feedback (VF) are weighted flexibly depending on the task and on the quality and availability of feedback (Sober and Sabes, 2003; 2005). In this perspective, externally altering the natural “gain” of proprioception through sub-sensorial stimulation could affect the sensory weighting that happens during the task and impact motor performance. It cannot be excluded also that the weighting process could completely “bypass” the artificial sensory boost.

In this work we aim at testing if enhancing proprioception through sub-sensorial electrical stimulation can alter motor performance during reach-and-hold and visuomotor adaptations (VMA) tasks with different VF conditions. As motor adaptation is considered one of the processes constituting motor learning (Shadmehr and Wise, 2005; Krakauer, 2009), our experiments aim also at giving additional information on the usability of SR stimulation as an additional aid during rehabilitation therapy of reaching movements. In our experiments, we asked subjects to perform a reach-and-hold task where VF was removed during the hold part of the task. Subjects repeated the task several times with different levels of sub-sensorial stimulation applied to the muscles driving the movement. This experiment was designed for determining the subject-specific optimal stimulation level, defined as the current level minimizing movement variability during the hold phase of the movement when VF was not present (thus in the portion of the task that was only reliant on proprioceptive feedback). Subjects were then split in three groups and each group performed a version of a visuomotor adaptation experiment twice, once with optimal sub-sensory stimulation (*Stim* condition) and once with no stimulation (*NoStim* condition), in a random order. One group performed the experiment with the VF always present (*Full VF*), one with VF limited to the initial part of the reaching movement (*Limited VF*) and one with VF only of the starting positions and end results of each movement (*No VF*). These three VF conditions were selected to examine the impact of enhancing proprioception in both the planning and on-line adjustment phases of the movement. We report here a limited effect of sub-sensory stimulation only when the VF is not present. These findings have major implications for evaluating the use of sub-sensory electrical stimulation during the execution of complex tasks.

## 2. Methods

### 2.1 Participants

A total of 40 healthy individuals (19 females, age 24.0 ± 4.3 years) volunteered for this study by signing an informed consent. All the experimental procedures were approved by the Ethical Committee of University College Dublin and have been conducted according to the WMA’s declaration of Helsinki. No personal or sensitive data were collected for the study. This consisted of four different experiments executed using the same experimental setup. Each subject participated to two of the experiments twice (one common to all subjects and one group-specific) during two experimental sessions performed in different days, often within the same week.

### 2.2 Experimental Setup

During all experiments, subjects sat on a chair placed in front of a computer screen placed at a distance of 1 m (**Fig. 1**). The elevation of the chair was controlled so to keep the shoulder abducted at 100°. Subjects had the right hand strapped to a manipulandum attached to a tri-axial load-cell (3A120, Interface, UK), while the wrist and the forearm were wrapped to the support plan and immobilized using self-adhesive tape. The elbow and shoulder flexion were fixed at 90° and 60°, respectively. All experiments consisted in the exertion of isometric forces against the manipulandum, as instructed by a virtual scene presented on the screen. The virtual scene consisted of a grey cursor, commanded in real time by the *x* and *y* components of the force exerted on the manipulandum, a filled circle indicating the center of the exercise space (0 N of force applied) and a target, represented by a hollow circle. The center and target circles had a radius of 0.7 cm or 1.2 cm, depending on the experiment (see 2.3 and 2.4). Targets were always placed at a distance from the center equal to 7.5 cm on the screen, equivalent to 12 N force exerted in the direction of the target. Data from the load-cell were sampled at 50 Hz. All the software constituting the virtual scene was custom developed in Labview.

**Figure 1.**
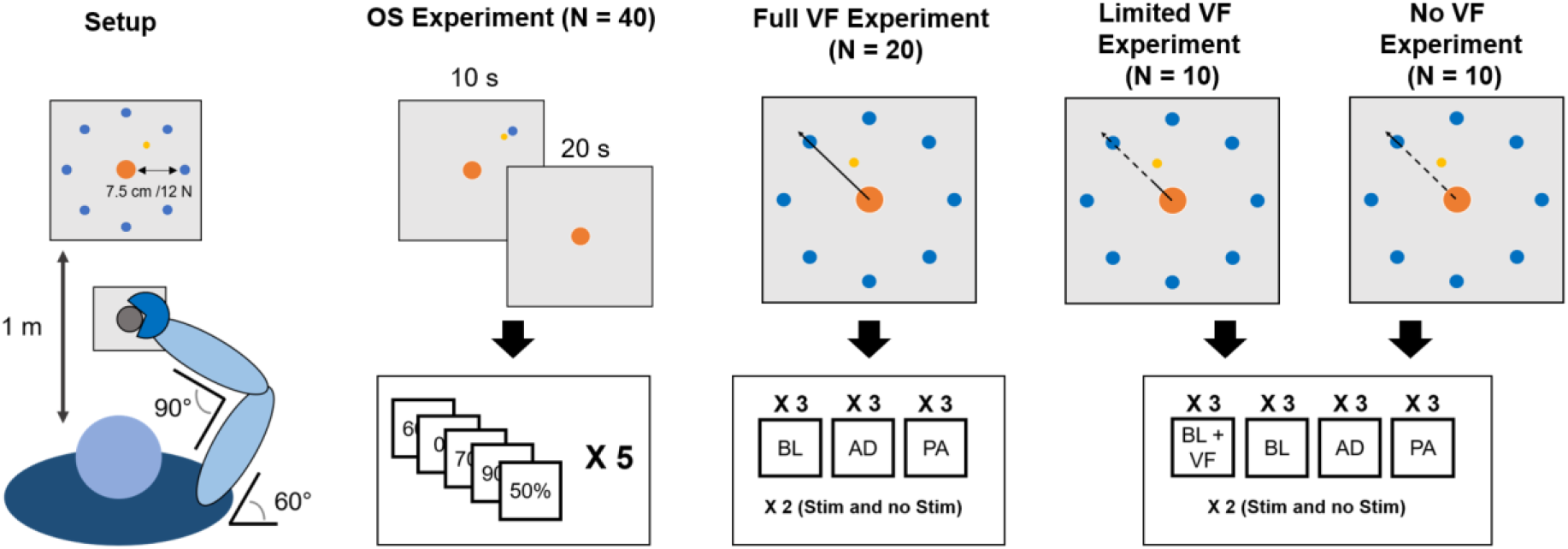
Experimental Setup. Subjects maintained the same position (leftmost panel) through all the experiments. During the OS experiment subjects had visual feedback during the reaching part of the trial and the first 10 seconds of holding and no visual feedback for the remaining 20 seconds. In the VMA experiments, visual feedback (bold line marks when it is present, dashed line when it is absent) changed across the different versions of the experiment. In the Full VF version feedback was always present. In the Limited VF version feedback was present only in a 2 cm radius from the center. In the No VF version feedback was present only for distances longer than that of the target.

### 2.3 Sensory Threshold Selection

At the beginning of each experimental session for each subject, a procedure for the identification of the subject- and session-specific sensory threshold (ST) was performed. Two electrodes for electrical stimulation (5×5 cm, Valutrode Lite, Axelgaard, US) were positioned on the lateral head of the triceps brachii (TLH) muscle. This muscle was chosen as in this type of setup has proven to be the muscle majorly involved in reaching the upper right part of the workspace (De Marchis et al., 2018). The electrodes were placed at about 2/3 the length of the muscle belly in each direction. The ST was defined as the smallest noise-stimulation current (white Gaussian noise, bandwidth 0.1-1000 Hz) that the subject could perceive and was calculated by iteratively increasing the root mean square value (RMS) of the stimulation noise by 10 μA (starting from 0) every 30 seconds until the subject started feeling a clear tingling sensation under the electrodes. Stimulation was administered using a voltage-driven current stimulator (Model 2200, A-M Systems, US), commanded using a custom software developed in Labview. The ST level was estimated for each subject during each experimental session.

### 2.4 Optimal stimulation experiment

The study consisted of four different experiments all executed using the setup just described. The experiments consisted of an optimal stimulation (OS) experiment (to which all subjects participated twice) and in three different versions of a visuomotor adaptation (VMA) experiment (of which each subject experienced only one version, twice), each version characterized by a different VF on the cursor trajectory presented during the task performance.

The aim of the OS experiment was to determine the session-specific optimal stimulation level for each subject, defined as the level of sub-sensory stimulation that maximizes performance by decreasing task variability in the absence of VF. During the OS experiment subjects performed a series of reach-and-hold tasks, consisting of reaching for a target of 0.7 of diameter positioned in the upper right side of the screen (**Fig. 1**) and of holding the cursor as close as possible to the center of the target for 30 seconds. The VF was projected on the screen only during the reaching phase and for the first 10 s after they reached the target, and was then removed. During each task, subjects received sub-sensory noise stimulation on their TLH muscle at six different current levels, equal to 0% (no current), 50%, 60%, 70%, 80% or 90% of their ST. Subjects experienced each level of sub-sensory stimulation five times in a random order, for a total of 30 repetitions (6 current levels × 5 times). The session-specific OS level was estimated at the end of each OS experiment as the percentage of ST (excluding 0% ST) yielding the smallest average (across the 5 repetitions for each percentage) standard deviation in the Cartesian distance between the cursor and the target during the 20 s of the hold phase of the task where the visual feedback was not present (*stdDist*). Additional analyses were performed in post processing. Specifically, we checked for statistically significant differences (Wilcoxon’s signed rank test, α=0.05) in the average *stdDist* between OS and 0% ST across all subjects. We then analyzed the distribution of the OS percentages across the different stimulation levels, for both OS experiments of all subjects. Finally, we analyzed the relationship between the stimulating current and the motor performance by fitting a first order polynomial, using a least square algorithm on the average *stdDist* values relative to each stimulation intensity. The quality and significance of the fitting was evaluated by calculating Pearson’s coefficient *ρ*.

### 2.4 Visuomotor adaptation experiment

All three versions of the VMA experiment consisted of isometric reaching movements where the subjects were asked to drive the cursor towards a random target (diameter 1.2 cm) presented at 7.5 cm (12 N) from the center. Subjects performed their assigned version of the VMA experiment immediately after the OS one, in both sessions.

The versions of the VMA experiment differed only in the VF that was provided to the subject during the reaching tasks. 20 subjects (9 females) performed the VMA experiment with continuous view of the movements of the cursor they were driving (*Full VF*). 10 subjects (2 females) performed the VMA experiment while receiving VF of the movement of the target only up to 2 cm (3.3 N) from the center of the virtual scene (*Limited VF*). Finally, 10 subjects (8 females) performed the experiment with no VF (*No VF*) on the movement of the cursor during the trajectory. In the *No VF* experiment subjects were shown the cursor only between 0 and 0.5 cm (0.7 N) from the center and after exceeding the distance to the center of the target (7.5 cm, 12 N). Thus, in the *No VF* experiment subject received feedback only on the result of their reaching trial.

The VMA experiment consisted of 9 blocks during which the VF condition was applied. In the first 3 blocks (baseline, BL1-BL3) participants were asked to reach to 8 targets positioned in a compass-like configuration for 5 times in a random order (Figure 1). During these and subsequent blocks they were instructed to reach for the targets as fast as possible and they were given positive feedback (consisting in the target becoming green) if they were able to reach for the target in less than 1.5 seconds, and negative feedback (consisting in the target becoming red) otherwise. The feedback on the speed of the trial indicated by the change in color of the target was present in all three VF conditions. The targets for which a subject received negative feedback were appended and repeated at the end of the block, thus making each block consisting of 40 movements plus the repetition of all the negative-feedback movements. After the BL blocks, subjects performed three adaptation blocks (AD1-AD3) where they were asked to reach for the targets while adapting to a 45° clockwise rotation applied to the mapping between the force sensor and the virtual scene. The only instruction that the subjects were given was to try to obtain positive feedbacks on their movements. Also in this case, subjects performed 5 repetitions of all 8 targets in a randomized order (40 tasks), and repeated the targets for which they received negative feedback at the end of the trial. Finally, subjects performed three unperturbed post-adaptation blocks (PA1-PA3) that were used to washout the adapted motor plan. Subjects performing the *Limited VF* and *No VF* VMA versions also experienced 3 additional blocks before the BL ones, that consisted of unperturbed baseline blocks with full VF (BL-VF). The aim of these blocks was to allow the subjects to practice and fully understand the task before the limitation to the VF was applied. Subjects performed their assigned VMA experiment in both experimental sessions, once while receiving sub-sensory stimulation (through all the 9 blocks of the experiment) at the OS level calculated in that same experimental session (*Stim*), and once without stimulation (*NoStim*), in a random order. The stimulation level used during the *Stim* condition was the one identified for the subject during the OS experiment of that specific session. Participants were blinded to the condition. For all three versions of the VMA experiment, half of the assigned subjects performed the *Stim* condition in the first experiment, the other half in the second experiment. For each reaching repetition, we analyzed the center-out portion of the movement, from the moment in which the cursor exited the origin target to the moment it reached the goal target. Each center-out movement was extracted and length-normalized over 100 data points. We analyzed the trajectory data by means of two metrics (**Fig. 2**): the initial angular error (IAE) and the normalized curvilinearity (NC). The IAE was calculated as the angle between the straight line connecting the ideal path and the actual path of the movement at 2 cm from the origin. This distance was selected because subjects performing the *Limited VF* experiment had the VF removed after 2 cm, thus for them this metric represents the angular error before losing VF. The NC was defined as the ratio between the actual distance covered by the cursor between the center and the target and the length of the straight line connecting the center and the target. The IAE is intended to capture the error in movement planning before the onset of potential compensations, while the NC metric accounts for both the initial movement error and the changes in motor plan that the subject undergoes to compensate for the shooting error. The analysis of IAE and NC was performed on the first 40 movements of each block (thus excluding the repeated trials in each blocks) and the behavior of the two metrics was analyzed both movement-by-movement and as average in each block. Moreover, the analysis were differentially performed on all targets together and by considering only the targets were the triceps are active (that are, using a compass notation, targets N, NE and E, as estimated in (De Marchis et al., 2018) using the same experimental setup) or the targets were the triceps are not involved (all targets excluding N, NE and E).

**Figure 2.**
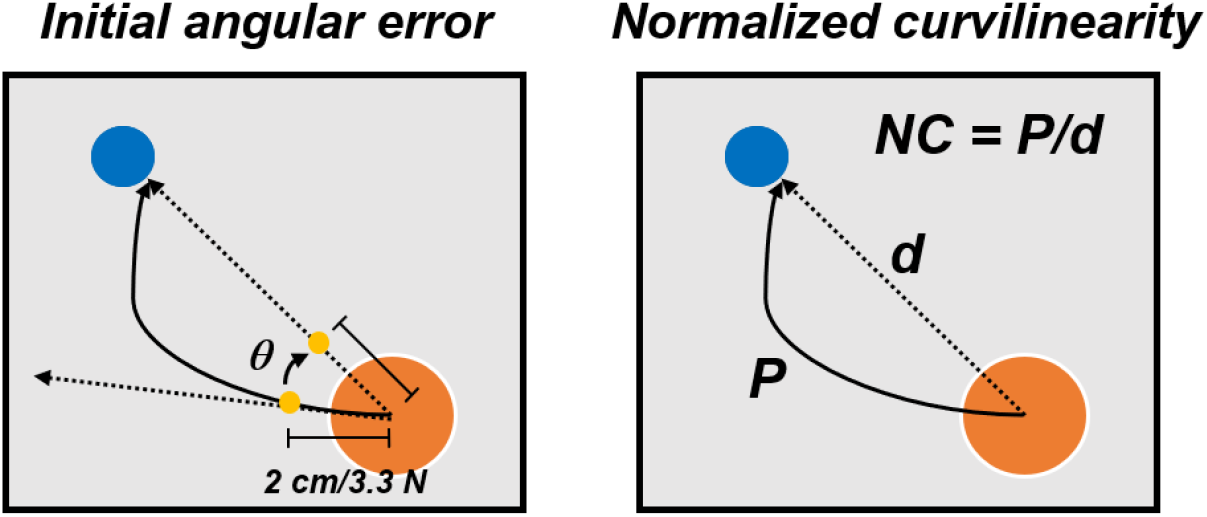
Performance metrics for the VMA experiment. The initial angular error (left) was calculated, for each movement repetition, as the angle between the actual and optima trajectories at 2 cm from the center of the workspace. The normalized curvilinearity (right) was calculated, for each movement repetition, as the ratio between the actual movement path and the ideal one.

Finally, for all three versions of the VMA experiment, we compared the number of repeated trials (thus the number of errors) across subjects in the first block of adaptation (AD1) between the *Stim* and *NoStim* conditions. This comparison was based on a Wilconxon’s signed rank test with significance level α = 0.05.

## 3. Results

### 3.1 OS Experiment

The results of the OS experiments performed by the subjects in the two experimental sessions were pooled together in the analysis. Thus, the 80 instances (40 subjects × 2 experimental sessions) were treated as independent measures. As expected from similar experiments (Magalhaes and Kohn, 2012; 2014; Severini and Delahunt, 2018), we consistently observed a decrease in accuracy during the hold-phase of the OS task when the VF was removed (example for one trial of one subject in **Fig. 3A**). From the analysis of the OS levels, considering also the trials where no current was actually applied (0%), we observed that in 7 instances out of 80, the average *stdDist* was lower for 0% stimulation than for a stimulation level above 0%. This accounts for 8.75% of the instances, against a value expected by chance of 13.33% (**Fig. 3B**). For the instances in which the 0% level presented the lowest average value of *stdDist* across the task repetitions, the value of OS was selected as the value of actual stimulation (thus above 0%) which yielded the lowest average *stdDist* (**Fig. 3B**). The OS levels were mostly distributed towards percentages close to the ST (**Fig. 3B**) with 59 out of 80 OS levels observed for percentages of ST above 70%. We observed statistically significant lower values of *stdDist* for OS with respect to 0% stimulation (p<0.01 using Wilcoxon’s signed rank test), also considering the instances were 0% yielded the average lower *stdDist* results (**Fig. 3C**). Finally, we analyzed the correlation between the RMS of the stimulation current and the *stdDist* metric.

**Figure 3.**
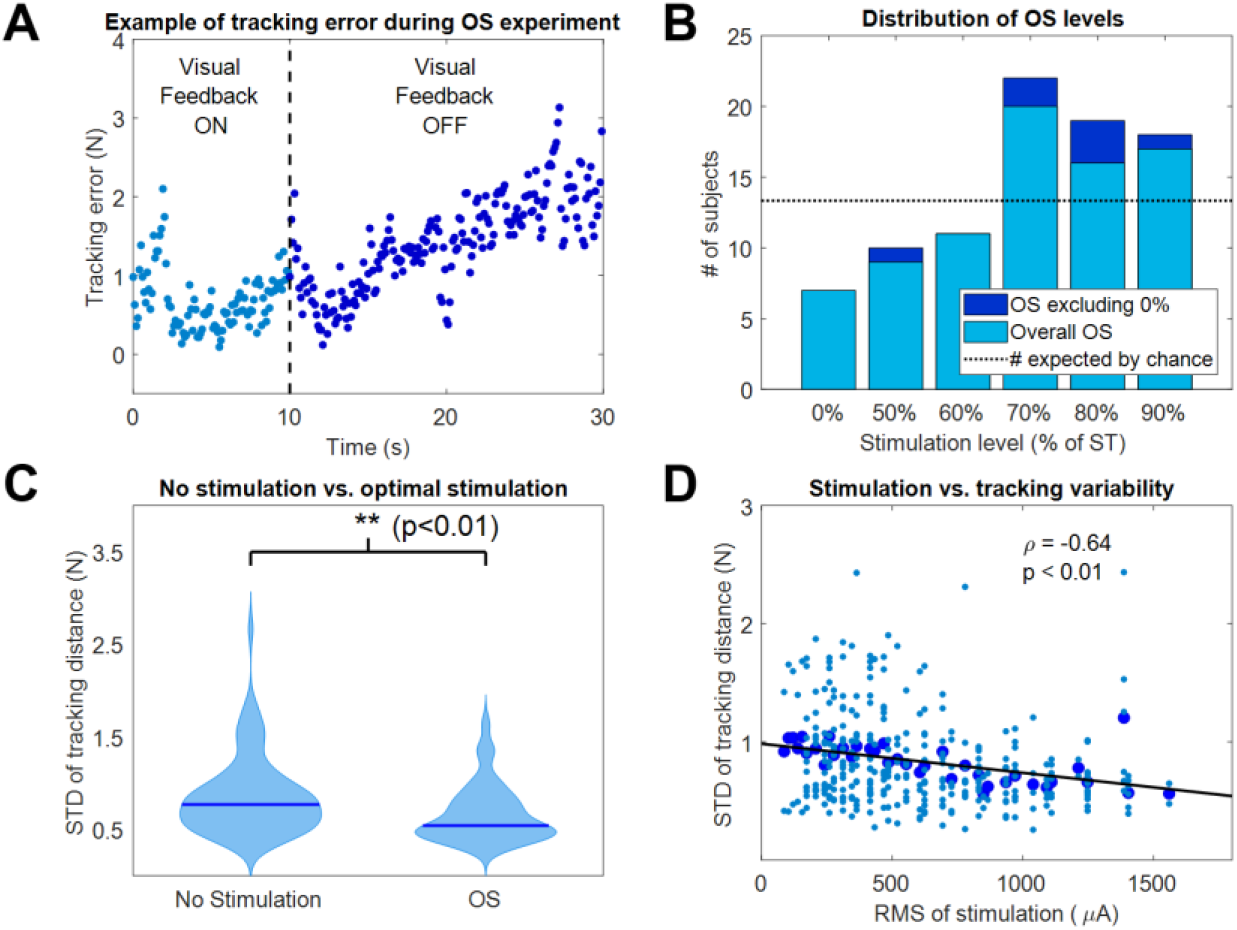
Results of the OS experiment. (A) Example of tracking error during a representative instance of the OS experiment. Movement variability around the target position increased as visual feedback was removed. (B) Distribution of the OS values, both including (light blue) and excluding (dark blue) the 0% level. (C) Violin plots of the tracking variability between OS values and 0% (no stimulation) values. ** indicates significant differences (Wilcoxon’s signed rank test) with p < 0.01. (D) Correlation between the RMS of stimulation and the STD of the tracking distance during the OS experiment.

We observed a negative correlation (**Fig. 3D**) between the average *stdDist* metric (averaged across all repetitions associated with a specific current level across subjects) and the relative RMS values of stimulation current, characterized by a significant (p<0.001) linear fitting with *ρ*=−0.64. This fitting indicates that, in our dataset, the performance increases with the RMS of the stimulation.

### 3.2 VMA Experiments

The results for the *Full VF* version of the VMA experiment (**Figure 4**) were in line with what had been observed in literature (Shadmehr and Mussa-Ivaldi, 1994) (**Fig. 4A**). Subjects presented marked movement errors, reflected in both the IAE and NC metrics, in the first block of perturbation (AD1) that were compensated over time. After-effects opposite to the direction of the original perturbation (in the IED) were present at the beginning of the post-adaptation phase and quickly vanished by the end of the experiment. When comparing the *Stim* and *NoStim* conditions, we were not able to observe substantial differences in trends in both metrics, such as a faster/slower adaptation speed or different values of IED or NC at the beginning of AD1 or at the end of AD3. This would have indicated, a higher/lower initial error and a higher/lower level of compensation of the error, respectively. Instead, both conditions presented remarkably similar trends in both metrics when considering all targets (**Fig. 4B**), only the targets where the triceps are active (**Fig. 4C**) and the targets where the triceps were not active (**Fig. 4D**). Similarly, we did not observe significant differences in the number of errors made by the subjects in AD1 between the two stimulation conditions for all the target groupings (rightmost panel, **Fig. 4B, 4C** and **4D**).

**Figure 4.**
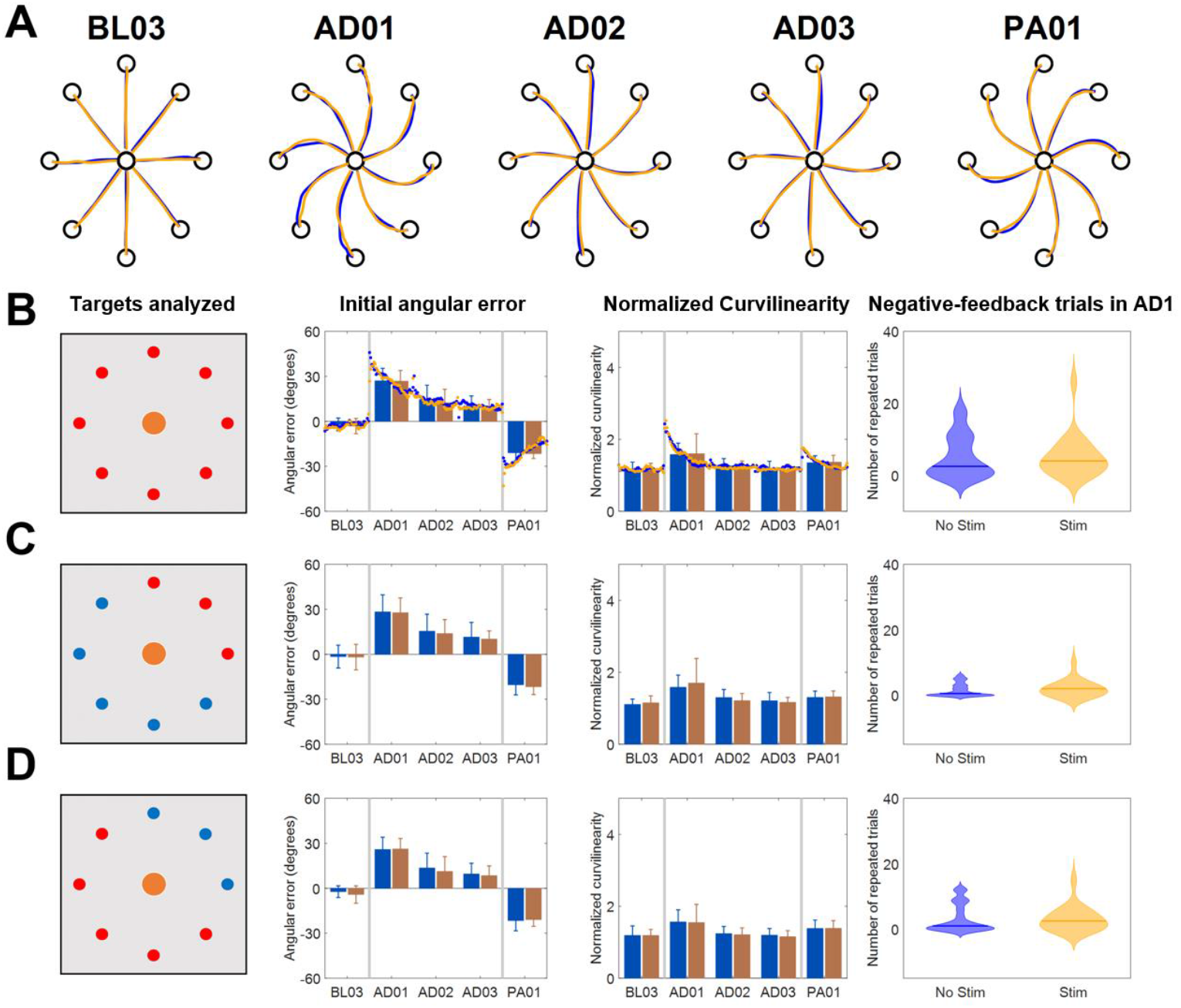
Results of the VMA *Full VF* experiment. (A) Example of force traces to targets for representatives blocks of the experiment. (B) Average, across subjects, performance metrics. The first panel from the left presents the targets analyzed (in red). The second panel presents the initial angular error metric, both as mean± standard deviation across the first 40 trials of each block (bars and whiskers) and as average (across subject) of the metric extracted for each single reaching movement for the first 40 trials (dots). The third panel presents the normalized curvilinearity metric, in the same notation. The fourth panel presents the violin plots of the number of negative-feedback trials (that had to be repeated) during AD1. (C) and (D) present the same results for only the upper right quadrant targets of the workspace (where the muscles stimulated are active) and for the remaining targets. In this case the metric plots are presented only as the mean± standard deviation across the trials of those targets in each block. In all plots, blue indicates the *NoStim* condition, Orange the *Stim* condition.

In the *Limited VF* version of the VMA experiment (**Figure 5**), trajectories were characterized by initial shooting errors followed by abrupt deviations once the VF was removed (**Fig. 5A**). As the AD blocks progressed, subject showed decreased shooting errors (also captured by a progressive decrease in IAE and NC) but still exhibited abrupt modifications in their trajectories once the feedback was removed. When comparing the *Stim* and *NoStim* conditions we observed a qualitative trend where *Stim* trials were characterized by higher initial values of IAE and NC at AD1 with respect to *NoStim*. The two conditions exhibited similar values on both metrics at AD3. The trends observed appeared to be present on all targets, regardless of groupings (**Fig. 5B, 5C** and **5D**). Finally, we did not observe significant difference in the number of errors at AD1 between the two conditions.

**Figure 5.**
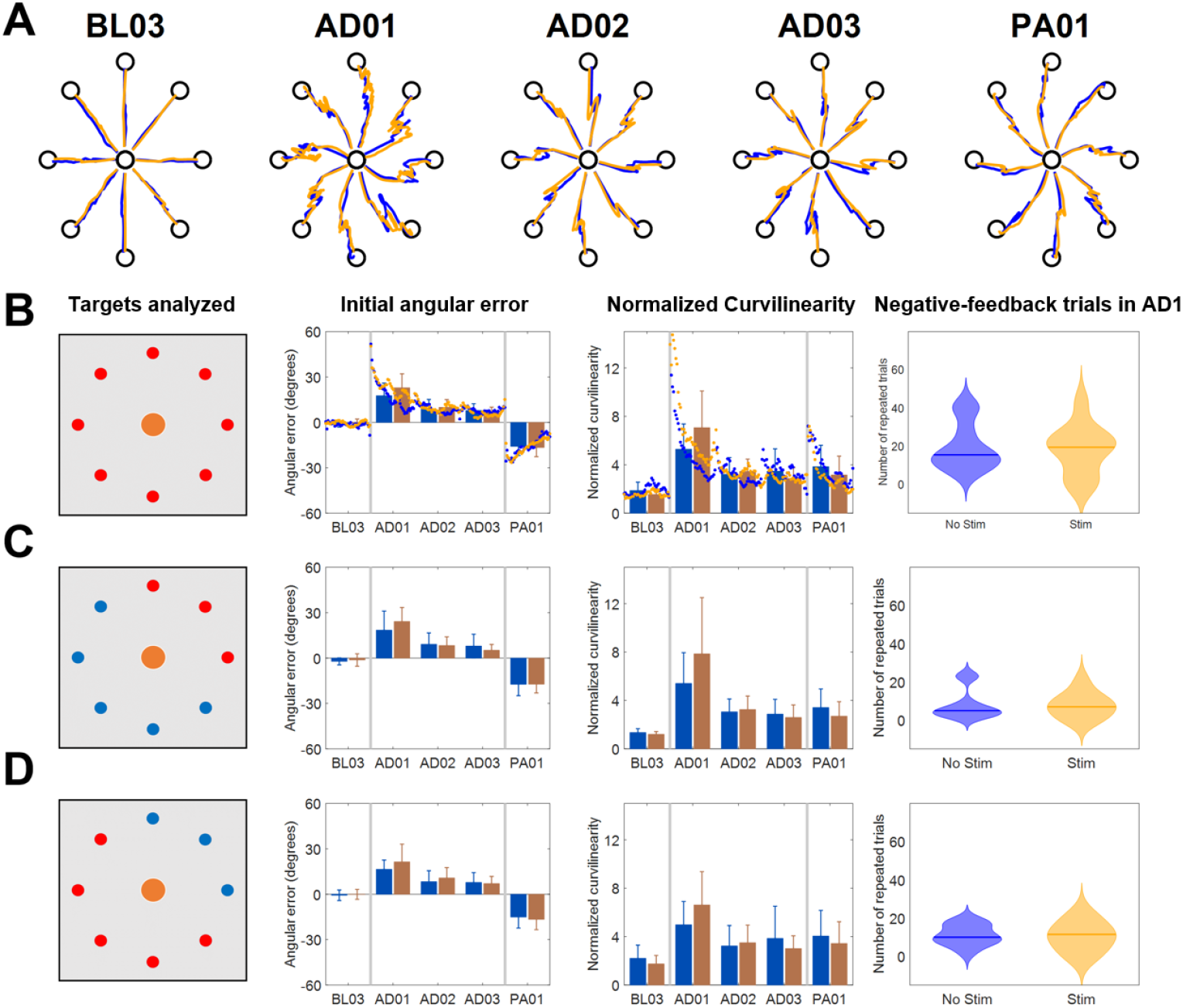
Results of the VMA *Limited VF* experiment. (A) Example of force traces to targets for representatives blocks of the experiment. (B) Average, across subjects, performance metrics. The first panel from the left presents the targets analyzed (in red). The second panel presents the initial angular error metric, both as mean± standard deviation across the first 40 trials of each block (bars and whiskers) and as average (across subject) of the metric extracted for each single reaching movement for the first 40 trials (dots). The third panel presents the normalized curvilinearity metric, in the same notation. The fourth panel presents the violin plots of the number of negative-feedback trials (that had to be repeated) during AD1. (C) and (D) present the same results for only the upper right quadrant targets of the workspace (where the muscles stimulated are active) and for the remaining targets. In this case the metric plots are presented only as the mean± standard deviation across the trials of those targets in each block. In all plots, blue indicates the *NoStim* condition, Orange the *Stim* condition.

In the *No VF* version of the VMA experiment (**Figure 6**), once again we observed initial changes in both metrics at AD1 due to the rotation. These changes were compensated over the trials even without VF (consistently with what shown in Scheidt et al., 2005) although to a smaller level with respect to the *Full VF* experiment (**Fig 6A** and **4A**). Also in this case, the adaptation behaviors were reflected in both metrics. We did not observe differences in the behavior of the IAE and NC metrics between the two stimulation conditions, either for all the targets or for the different groupings. However, the *NoStim* condition presented a significant higher number of reaching errors at AD1 with respect to the *Stim* condition that was observed for all the targets togethers (p = 0.046, **Fig. 6B**) and for the grouping representing only the targets were the triceps were active (p = 0.043, **Fig. 6C**).

**Figure 6.**
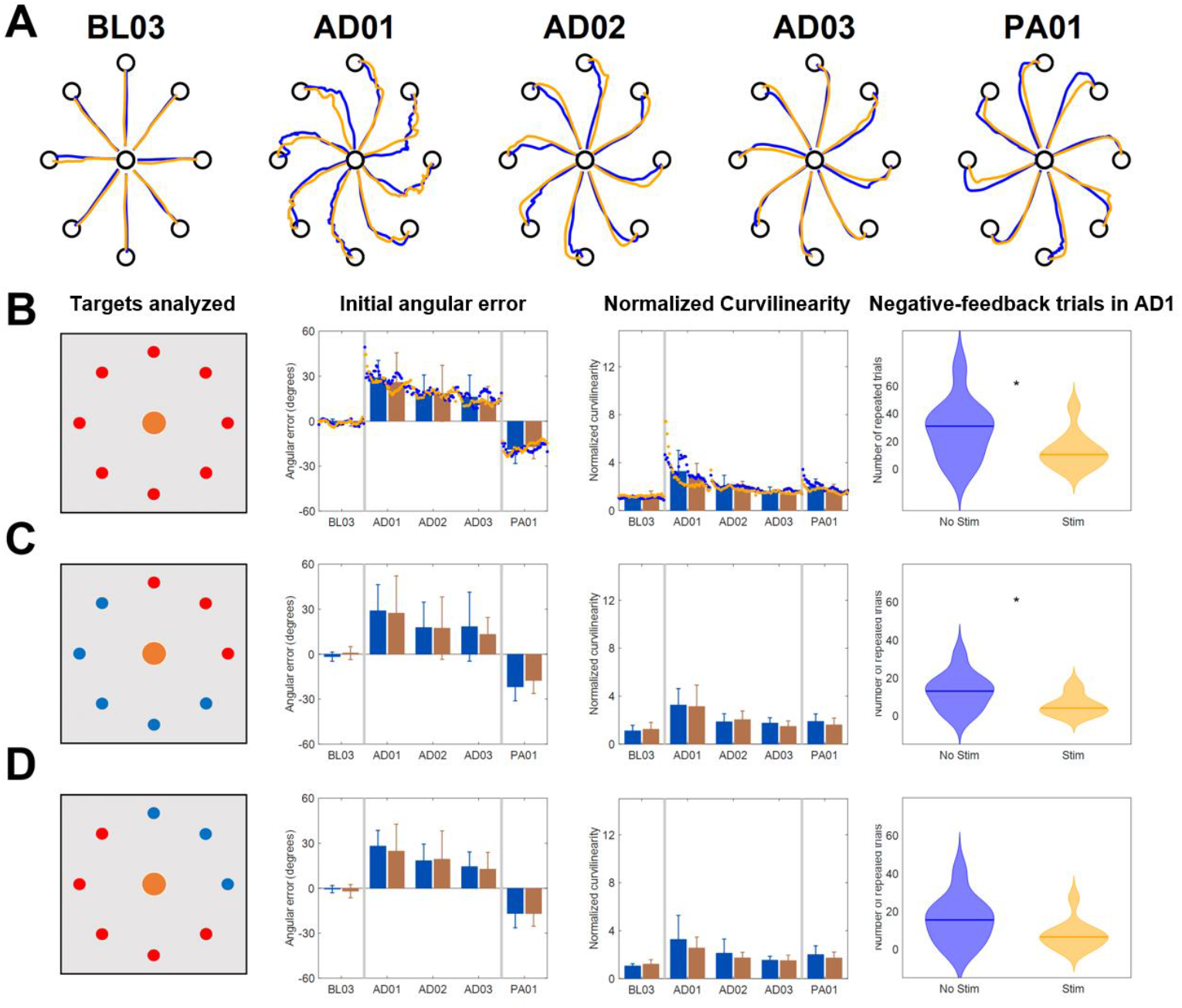
Results of the VMA *No VF* experiment. (A) Example of force traces to targets for representatives blocks of the experiment. (B) Average, across subjects, performance metrics. The first panel from the left presents the targets analyzed (in red). The second panel presents the initial angular error metric, both as mean± standard deviation across the first 40 trials of each block (bars and whiskers) and as average (across subject) of the metric extracted for each single reaching movement for the first 40 trials (dots). The third panel presents the normalized curvilinearity metric, in the same notation. The fourth panel presents the violin plots of the number of negative-feedback trials (that had to be repeated) during AD1. * indicates significant differences (Wilcoxon’s signed rank test) in the number of negative-feedback trials with p<0.05 (p = 0.046 in (B) and p = 0.043 in (C)). (C) and (D) present the same results for only the upper right quadrant targets of the workspace (where the muscles stimulated are active) and for the remaining targets. In this case the metric plots are presented only as the mean± standard deviation across the trials of those targets in each block. In all plots, blue indicates the *NoStim* condition, Orange the *Stim* condition.

## 4. Discussion

In our results we observed that sub-sensory electrical stimulation was associated with smaller movement variability during the phase of the OS experiment where VF was not available and task performance depended solely on proprioceptive feedback. Moreover, we observed a correlation between stimulation current and movement variability whereas higher current levels were associated with better task performance across subjects. These results, taken together, further confirm that sub-sensory electrical stimulation can improve task performance in tasks were proprioception is the primary feedback modality (Gravelle et al., 2002; Ross and Guskiewicz, 2006; Collins et al., 2014; Severini and Delahunt, 2018).

On the other hand, we observed only small evidence of an effect of the stimulation during the VMA experiments, that was mainly characterized by a significant decrease in negative-feedback movements (that are movements that took more than 1.5 seconds for the subject to complete) between the two stimulation conditions during the first block of adaptation for the subjects that performed the *No VF* version of the experiment. When the VF of the trajectory was present, completely or partially, we did not observe substantial differences in task performance, as captured by two different metrics, between the *Stim* and *NoStim* conditions other than a qualitative slight decrease in task performance during AD1 for the *Limited VF* group. In the following, we will further discuss upon these results.

The results of the OS experiment provide, in this study, the strongest evidence of the effectiveness of sub-sensory stimulation in boosting proprioception and influence task performance. In the OS experiment we did not observe a clear SR behavior, characterized by a U-shaped relationship between the change in performance and the intensity of the stimulation (Collins et al., 1995). Such behavior is unlikely to appear in a group analysis (Bates, 1996; Severini and Delahunt, 2018), given the differences in ST across subjects and across different sessions for the same subjects that have been observed in this and previous studies (Magalhaes and Kohn, 2012; 2014). Nevertheless, we did observe a significant negative correlation between the stimulation intensity and the tracking error (**Fig. 3D**), suggesting that sub-sensory stimulation is more effective as its intensity increases. This linear relationship does not rule out the presence of a SR-like behavior, but hints that such behavior may arise by considering stimulation intensities that are above the ST of subjects. On the other hand, stimulating currents above ST could lead to additional confounding factors affecting motor task performance related to the increase in attention or arousal, and the few studies that investigated the use of supra-sensory stimulation levels in humans found that it leads to an overall decrease in performance (Iliopoulos et al., 2014; Severini and Delahunt, 2018). The results of the OS experiment support the design choice of using sub-sensory stimulation levels close but below ST (frequently 90% of ST) that is often employed in similar studies (Gravelle et al., 2002; Magalhaes and Kohn, 2012; 2014).

In contrast with the results obtained in the OS experiment, we observed little evidence of an effect of the stimulation during the different VMA experiments. In the *Full VF* version of the experiment, the adaptation patterns were remarkably similar between the two stimulation conditions. We observed some small differences in performance between the two stimulation conditions in the first block of adaptation for both the *Limited VF* and the *No VF* versions of the experiment, although results appear to be more solid in the latter rather than the former. In the *Limited VF* experiment we qualitatively observed higher values in both performance metrics during AD1 for the *Stim* condition. In the *No VF* experiment we did not observe differences in trends between the two metrics, but the *Stim* condition was characterized by a significant smaller number of negative-feedback trials, especially for the targets of the upper right quadrant, where the muscle undergoing stimulation was active. Both the trends that we observed in the *Limited VF* and *No VF* experiments could be potentially explained by the stimulation impacting the weighting process between proprioceptive and visual feedbacks that happens during reaching tasks in general, and motor adaptations in particular. Previous studies have shown that different feedback modalities mix flexibly during the execution of voluntary movements and during motor adaptations (Sober and Sabes, 2003; Scheidt et al., 2005; Sober and Sabes, 2005; Shabbott and Sainburg, 2010). While visual feedback is responsible for estimating the limb position required in the planning of the movement trajectory, proprioception contributes in generating the necessary feedforward commands required for movement execution (van Beers et al., 2002; Sober and Sabes, 2003; 2005). Primary and secondary muscle spindles have been observed to increase their firing rates during isometric contractions (Edin and Vallbo, 1990), indicating that these afferents encode information on muscular state even if the muscles are not changing in length. A previous study on spindles behavior during visuomotor adaptations has shown that adaptation leads to a progressive decrease in the activity of the spindles (Jones et al., 2001). The authors linked this result to the fact that adaptation to visuomotor rotations is achieved by updating the internal models mapping the kinematics of the movement, a process relying mostly on visual and less on proprioceptive feedback (Krakauer et al., 1999; Krakauer, 2009), as confirmed also in a study involving individuals with proprioceptive deficits (Lajoie et al., 1992). Decreasing the weight of the spindles’ information during visuomotor adaptation would then help resolving the conflict between the visual and proprioceptive maps that the perturbation induces (Jones et al., 2001). This re-weighting of proprioceptive information has been shown to happen centrally, at the level of the somatosensory cortex, rather than at the spinal level (Bernier et al., 2009), and to be more prominent at the beginning of the adaptation period and then alleviated as the adaptation converges.

Thus, as the activity of the spindles is down-regulated at the beginning of adaptation, the supposed enhancement of such activity by the stimulation would effectively clash with the sensory re-weighting process. This clash, in the *Limited VF* experiment, where VF of the shooting error is provided but proprioceptive feedback is still necessary for successfully completing the task, could translate in bigger initial errors as the stimulation supposedly antagonizes the spindle down-regulation. The fact that a similar effect is not present if the *Full VF* experiment could be explained by the primacy of VF over proprioception during visuomotor adaptations that bypasses the potential effects of the stimulation. On the other hand, in the *No VF* experiment, where proprioception is the only available feedback modality, its supposed boost through the stimulation may lead to increased feedback reliability that may translate in a smaller number of negative-feedback trials.

These explanations, although plausible, cannot be fully confirmed from these results due to the small sample examined in the experiments from which they have been derived, that must be listed as main limitation for the study herein presented. Another potential limitation of this study, that could also help explain the differences in stimulation effectiveness that we observed between the OS and VMA experiments, could be represented by the fact that we selected the optimal stimulation level based on the performance during the holding phase of the OS experiment and then tested it during a reaching task in the VMA experiments. In a literature review recently published by Shadmehr (Shadmehr, 2017) the author raised the possibility that these two tasks (holding and reaching), similarly to what happens during ocular movements, employ different neural circuitries. In this interpretation, the discrepancy in stimulation effectiveness that we observe could be explained by an experimental design flaw where we used optimal currents derived from the holding task in a task that employs different neural circuits. Nevertheless, although there is evidence on the different nature of neural inputs during reaching and holding, no information is available on if and how proprioceptive feedback is processed differently between these two tasks.

To summarise, the results presented in this study further support the hypothesis that sub-sensory currents applied to the surface of the muscles affect proprioceptive feedback during movement, but this effect appears to be clearly beneficial for task performance only in tasks where proprioception is the primary feedback modality.

